# Retrieval induced forgetting in a non-monotonic hippocampal model

**DOI:** 10.1101/2023.08.25.544249

**Authors:** Benjamin Midler, James McClelland

## Abstract

Retrieval induced forgetting (RIF) occurs when the retrieval of one item negatively impacts the recall probability of related items stored in memory (Anderson et al., 1994). Recently, Ritvo et al. (2023) demonstrated RIF emerges in a neural network model equipped with non-monotonic plasticity. Their finding supports the non-monotonic plasticity hypothesis (NMPH; Ritvo et al., 2019): the theory that connection changes in the brain follow a “U” shaped function of post-synaptic stimulation. Here, we apply a unique implementation of non-monotonic plasticity to a neural network model of an idealized hippocampus (HPC) and evaluate it with an adaptation of a classic RIF task. The model evidences the behavioral and representational characteristics of RIF, replicating Ritvo et al. (2023). As a monotonic baseline model failed these tests, we provide evidence of non-monotonic plasticity’s sufficiency for RIF. In addition to demonstrating the NMPH is robust to multiple implementations and evaluative paradigms, we conduct additional analysis to provide a mechanistic explanation for how non-monotonic plasticity brings about RIF. Lastly, we evaluate the model with an expansion of RIF: reverse RIF. The model fails this final test, raising questions for future research on the necessary parameters of non-monotonic plasticity and whether it must pair with complementary processes in the brain.

## Introduction

Representations in memory dynamically interact. For instance, retrieving an item from memory inhibits recall of related memories and differentiates their representations. This phenomenon is called retrieval induced forgetting (RIF; Fig. 1). RIF is a robust effect demonstrated in both human behavioral and neuroimaging studies (Anderson et al., 1994; Wimber et al., 2015). A leading theory of the mechanism behind RIF is the non-monotonic plasticity hypothesis (NMPH; Ritvo et al., 2019). The NMPH holds that RIF is an emergent property of non-monotonic plasticity in the brain. Specifically, when the post-synaptic neuron receives negligible stimulation, its connection strengths to pre-synaptic neurons are negligibly altered. By contrast, moderate stimulation precipitates a downward turning of the connection strengths and high stimulation an upward tuning. This manifests as the characteristic “U” shaped plasticity function central to the NMPH.

**Figure 1:**
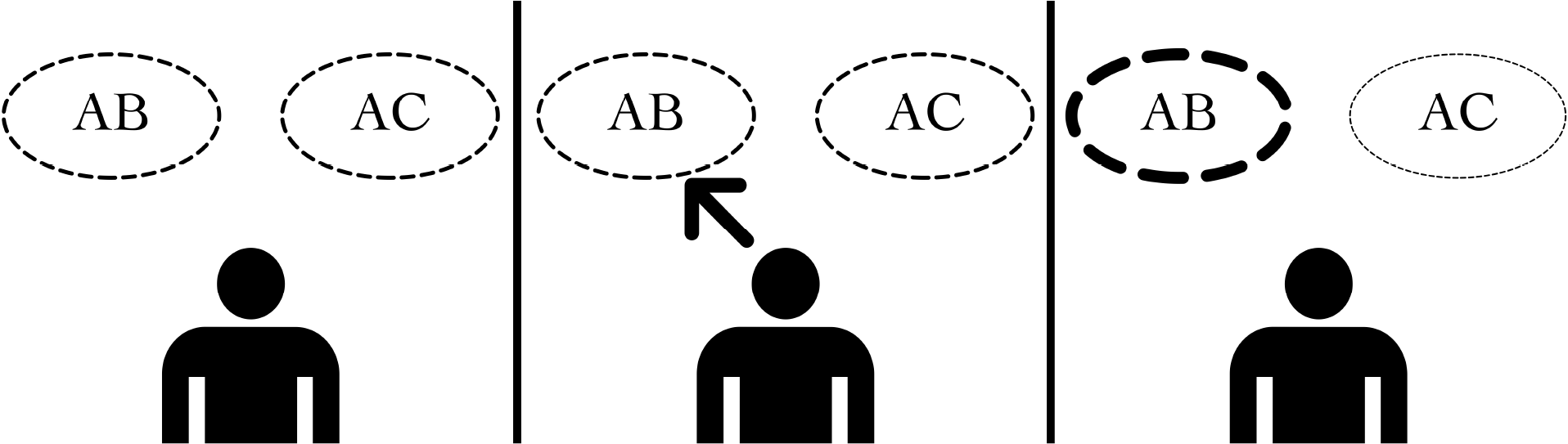
Retrieval induced forgetting. Retrieval induced forgetting (RIF) manifests when the retrieval of one item in memory (AB) not only strengthens its own memory trace and improves recall probability (indicated by thicker dashed line), but weakens the memory trace and decreases recall probability for a related item (AC), indicated by a thinner dashed line. RIF is often tested using language in a paired-associates paradigm and is a useful test of inter-representational dynamics in memory.

This process features in multiple theoretical accounts of neuroplasticity (Bienenstock et al., 1982; Norman et al., 2006), and may manifest biologically as homosynaptic LTP/LTD (Mulkey & Malenka, 1992; Hansel et al., 1996). Recently, Ritvo et al. (2023) constructed a neural network model of non-monotonic plasticity and demonstrated its ability to capture patterns of representational differentiation and integration from several experiments with human subjects, including an RIF task derived from Favila et al. (2016). Their work strengthens the argument that non-monotonic plasticity is the neural mechanism behind RIF and provides novel predictions, such as representational change being applied asymmetrically to items in memory.

Their four layer model makes use of a U-shaped plasticity function and inhibitory oscillations to instantiate non-monotonic plasticity—an approach previously emphasized as having a potentially key role in competition-dependent learning (Norman et al., 2006; Norman et al., 2007). In both this recent and prior works by the same group, the Bienenstock-Cooper-Munro (BCM; Bienenstock et al., 1982) learning rule was cited as an algorithm that likewise captures the NMPH and may give rise to the same neural dynamics (Norman et al., 2007; Ritvo et al., 2023). Here, we sought to test that conjecture. We developed a Hopfield network-based model of an idealized hippocampus (HPC) and equipped it with a learning algorithm derived closely from BCM. We evaluated the model with a unique adaptation of a classic RIF task, finding that the model produced key behavioral and representational characteristics of RIF, replicating the work of Ritvo et al. (2023) and proving the NMPH is robust to multiple unique implementations and evaluative paradigms. We likewise replicated their novel prediction of asymmetrical representational shift and conducted additional analyses to provide a mechanistic explanation for this effect. Finally, we tested the model’s ability to evidence reverse RIF—an expansion upon RIF involving re-learning forgotten items. Our model failed to capture the behavioral predictions of human subjects experiments. Therefore, while our model captured core characteristics of RIF, replicating Ritvo et al. (2023), it did not extend to a fuller suite of RIF behaviors. In sum, this work supports non-monotonic plasticity’s sufficiency for RIF and provides a mechanistic argument to that end.

## Methods

Computational modeling presents the opportunity to selectively and precisely control, measure, and manipulate the parameters of distinct neural processes. Fully-connected neural network models of an idealized HPC have yielded notable discoveries and proven themselves robust vehicles for studying mnemonic phenomena (e.g. Benna & Fusi, 2021). Thus, such models present an apt avenue to study the constituent processes of RIF, one that can be explicitly designed to instantiate a target subset of those processes. To that end, we adapted a connectionist model of an idealized HPC to instantiate the NMPH and tested it with an RIF paradigm adapted from the human subjects literature. The objective is to asses the model’s ability to capture key characteristics of RIF and to proffer a mechanistic explanation for that ability.

### Model design

Hopfield networks are connectionist models that are one of the earliest neural networks and have previously been used to model HPC processes (Hopfield, 1982; Krotov & Hopfield, 2020; Fig. 2). They, along with close derivatives, have been used to evaluate memory phenomena such as place cells and catastrophic interference (Hattori, 2010; Zhang et al., 2016; Benna & Fusi, 2021; Midler & McClelland, 2022). Furthermore, Hopfield networks can posses several qualities that lend themselves to modeling HPC, such as sparsity and connecting a smaller input layer to a larger hidden layer—features of both HPC itself and the input of cortical representations from entorhinal cortex (ERC) to HPC (Lipton & Eichenbaum, 2008; Lodge & Bischofberger, 2019).

**Figure 2:**
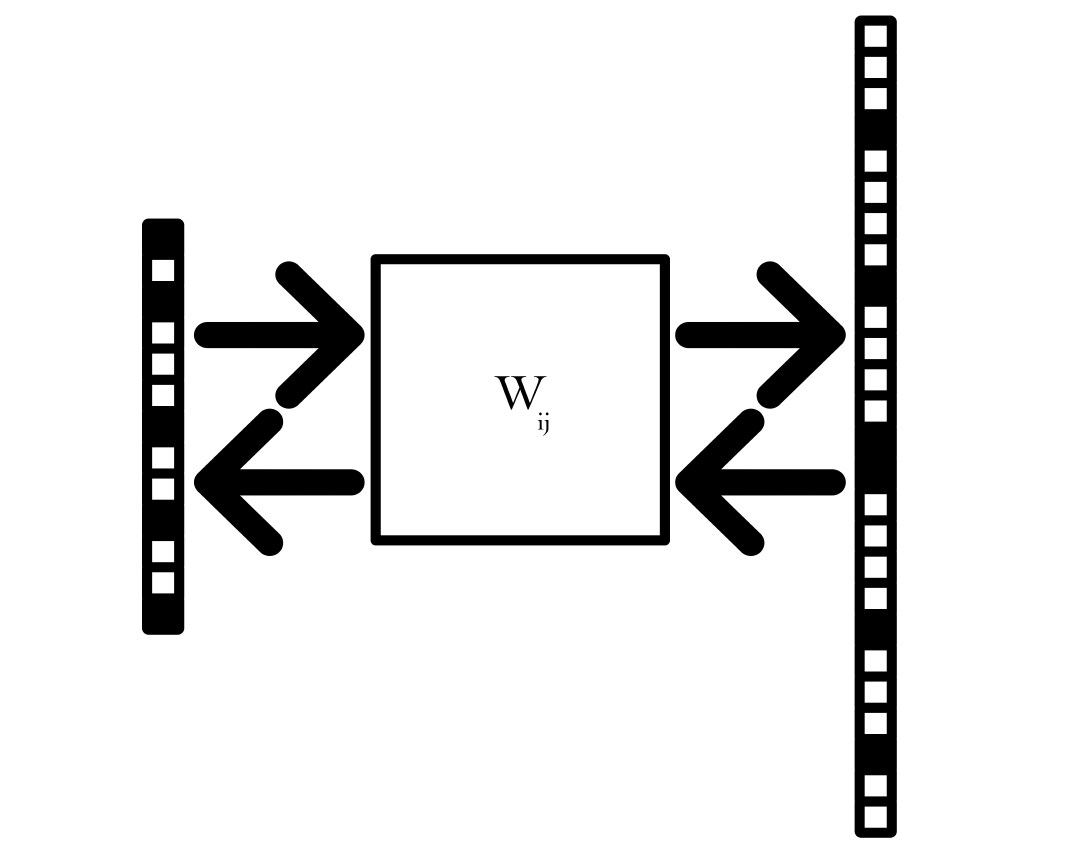
Hopfield network. Hopfield networks are connectionist neural networks that feed input patterns through a weight matrix (W_ij_) to form a hidden state (left to right). The hidden state then passes back through the transposed weight matrix to form a reconstructed pattern (right to left). These networks are models of an idealized HPC and are enhanced here with biologically plausible modifications, such as a k-winner non-linearity at the hidden layer and sparse connectivity.

We employ a modified modern Hopfield network that combines these structural similarities to HPC with further biologically plausible enhancements (Krotov & Hopfield, 2020). The model, dubbed the k-winner Hopfield network, applies a k-winner non-linearity to the hidden layer, forcing it to adopt distributed representations (Bhandarkar & McClelland, 2023). This is consistent with connectionist principles and does not sacrifice the HPC’s characteristic sparsity (O’Reilly & McClelland, 1994). Additionally, our model is adapted such that each hidden unit has reciprocal connections with 50% of units in the input/output layer. As each HPC neuron does not receive input from the entirety of ERC, this modification is consistent with the biology. This biologically plausible evolution of the modern Hopfield network thus incorporates several features that make it a suitable analogy to an idealized HPC and therefore a suitable testbed for the NMPH’s role in RIF.

### Learning rule

To instantiate non-monotonic plasticity in the k-winner Hopfield network, we adapted the BCM learning rule (Bienenstock et al., 1982). The change applied to the weight vector for the *i*th hidden unit, **Δw**_**i**_, is calculated by first taking the difference between the input vector,, and the weight vector for a given hidden unit, **w**_**i**_.

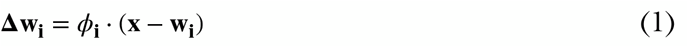

This difference is then scaled by *ϕ*_*i*_—the non-monotonic plasticity term set by post-synaptic stimulation. *ϕ* is similar to the non-linear threshold of the BCM algorithm (Bienenstock et al., 1982). However, *ϕ* is given by a non-monotonic function of current input to the post-synaptic unit rather than time-averaged activity (Dudek & Bear, 1992; Kirkwood et al., 1996). Specifically, *ϕ* is calculated by a quadratic function of each hidden unit’s percentile ranked net input, **a**_**i**_, and a fixed threshold, *μ*, dividing the positive and negative domains of the non-monotonic function, here set to 0.6, i.e. the 60th percentile.

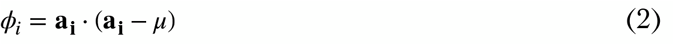

If **a**_**i**_ < *μ*, then *ϕ*_*i*_ is negative, causing **w**_**i**_ to be tuned away from **x**. If **a**_**i**_ > *μ* then *ϕ*_*i*_ is positive and **w**_**i**_ is tuned towards **x**. The function is therefore a manifestation of the “U” shaped relationship between activation and weight update described in the NMPH and employed by Ritvo et al. (2023; Fig. 3a). Featuring a fixed threshold between positive and negative domains, *μ*, may appear to be a departure from both the BCM algorithm—which employs a dynamic threshold set by time-averaged activity—and neurobiological evidence. However, as **a**_**i**_ is percentile ranked—not absolute—post-synaptic stimulation, *μ* is a variable quantity: it refers to a variable degree of stimulation (Kirkwood et al., 1996). In effect, the threshold employed here is dynamically set by averaged layer-wise activity at a single time point rather than by time-averaged activity at a single unit in BCM. *ϕ* either facilitates the weights tuning towards the input or away, depending on the level of post-synaptic stimulation, thus constituting non-monotonic plasticity. This learning rule is closely derived from BCM and endows the k-winner Hopfield network with the non-monotonic plasticity that should produce RIF, per the NMPH’s predictions.

**Figure 3:**
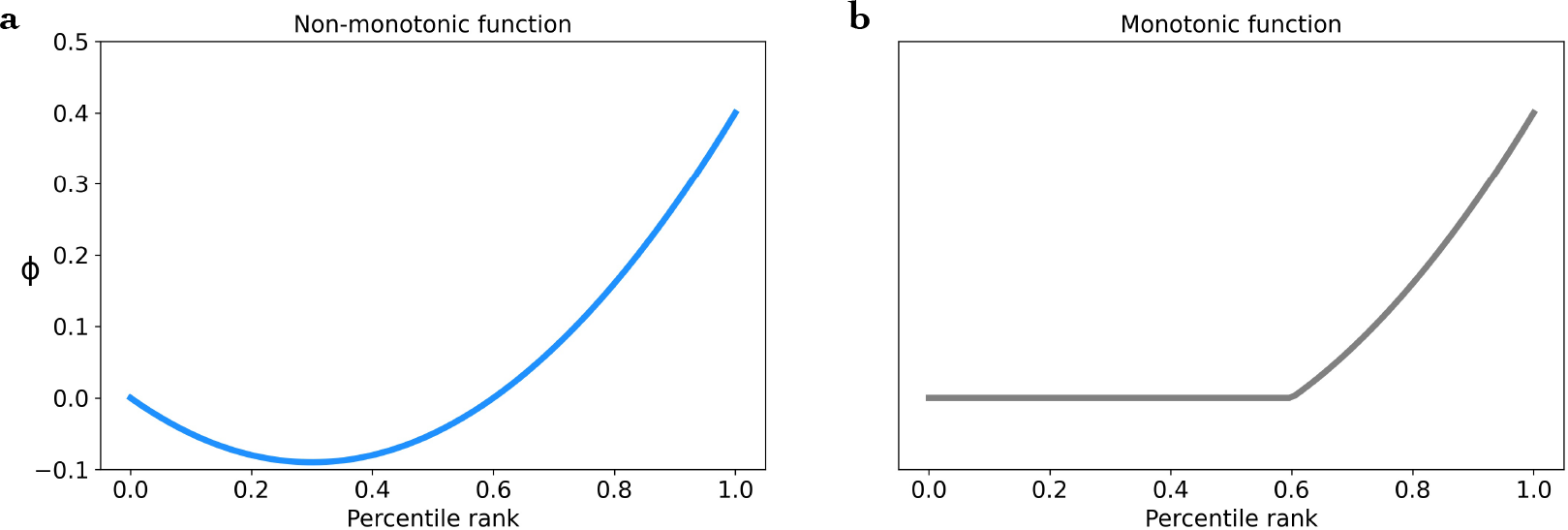
Plasticity functions. In order to implement non-monotonic plasticity, the plasticity function includes a scalar, *ϕ*; a function of the percentile ranked stimulation each hidden unit receives from the input pattern, **x**. The non-monotonic function (**a**) is the “U” shape characteristic of the NMPH. Thus, hidden units with moderate input will fall in the negative domain of the function (percentile rank <0.6), producing a negative *ϕ* and tuning the weights away from the input. By contrast, highly active units (percentile rank >0.6) will produce a positive *ϕ*, tuning their weights towards the input. The baseline monotonic function (**b**), is identical except, to make it monotonic, is bound to 0.

As a baseline, a second model was also evaluated. The baseline model was identical to the non-monotonic model in all but one respect; whereas the experimental model makes use of the non-monotonic plasticity function to produce *ϕ*, the monotonic baseline employs the same function with a lower bound of zero (Fig. 3b). The baseline model therefore employs strictly monotonic plasticity: its weights are either tuned towards the input or not at all; its weights are never tuned away. In sum, we applied a close adaptation of BCM to incorporate non-monotonic plasticity in a simplified model of HPC, constituting the properties theorized by the NMPH to be sufficient for RIF. This conjecture will be evaluated with an otherwise identical monotonic model as a baseline.

### Experimental task

The models’ ability to capture RIF was evaluated with a paradigm derived from Anderson and colleagues (1994). They described several experiments constituting one of the earliest RIF findings in humans. Their first experiment was adapted for use in this modeling context (Fig. 4).

**Figure 4:**
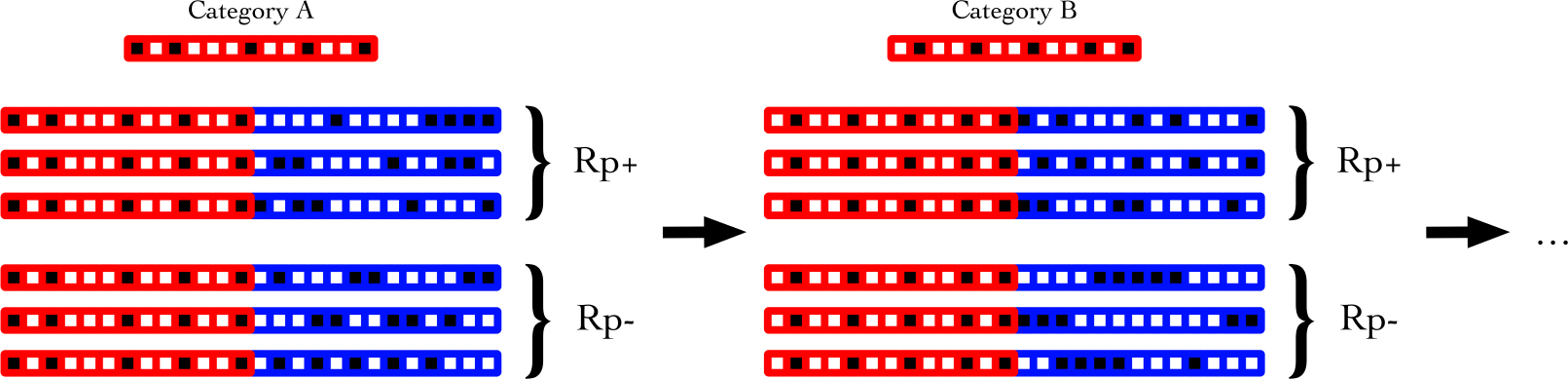
RIF paradigm. Inspired by Anderson and colleagues (1994), the model RIF task is a vector-based pair paradigm. For each of eight categories, a 50 unit long binary category vector with sparsity fixed to 10% is randomly generated (red). Six unique item vectors are then generated with the same parameters (blue), and concatenated with the category vector to form six unique pairs for each category. Three pairs are then randomly chosen to be re-encoded in the RIF task (Rp+), with the remaining three receiving no further practice (Rp-).

In the original experiment, eight category terms (e.g. fruit) were linked with six items in a paired associates memory task (e.g. fruit-apple). Of the six pairs in each category, three would receive additional retrieval practice, designated Rp+ pairs. The remaining three pairs in the category would receive no practice after the initial study phase, designated Rp-. Two categories out of the eight were held-out altogether, receiving no retrieval practice and designated Nrp. The hypothesis was that the Rp-pairs, as they were moderately similar to the Rp+ pairs due to their shared category term, would be subjected to more interference than Nrp pairs, constituting RIF.

For this modeling context, the paired associates paradigm made use of randomly generated binary vectors instead of words. For each category, a 50 unit long binary vector with sparsity fixed at 10% would be generated. Then, a further six item vectors with the same parameters would be generated and concatenated to the category vector, forming the six pairs of each category. The pairs would then be randomly split into Rp+ and Rp-groups. Although the modeling version of this RIF task preserves the Nrp categories, making an effective comparison between Rp pair performance and Nrp pair performance requires contextual information on the relationship between category and item. For instance, while human subjects know the item apple pairs with the category fruit and not the category furniture, the Hopfield network has no means to make such judgements; the relationship between category and item vectors is arbitrary. As such, Nrp performance is not evaluated, with focus instead paid to the contrast between Rp+ and Rp-performance and the experimental and control models.

Human subjects’ memory performance was evaluated by presenting a category cue and having subjects write down all items they recall paired with that category. In the modeling context, a category vector is passed into the model, producing a reconstruction. Cosine similarities between the reconstruction and all 48 patterns in the dataset are computed. The six most similar pairs are considered recalled if their similarity exceeds a threshold of one standard deviation above the mean. The proportion of correctly recalled items is then averaged across all categories for Rp+ and Rp-pairs and reported as the behavioral performance metric. Model weights are not updated during evaluation. The task itself is divided into three time points (Fig. 5). The first time point is at model initialization before any learning has occurred. The second time point is after the models have been trained on the entire dataset, and the third time point is after Rp+ pairs have been re-encoded. Thus, the experimental and control models are evaluated using an experimental assessment of RIF that has been validated with human subjects and is adapted to suit the modeling context.

**Figure 5:**
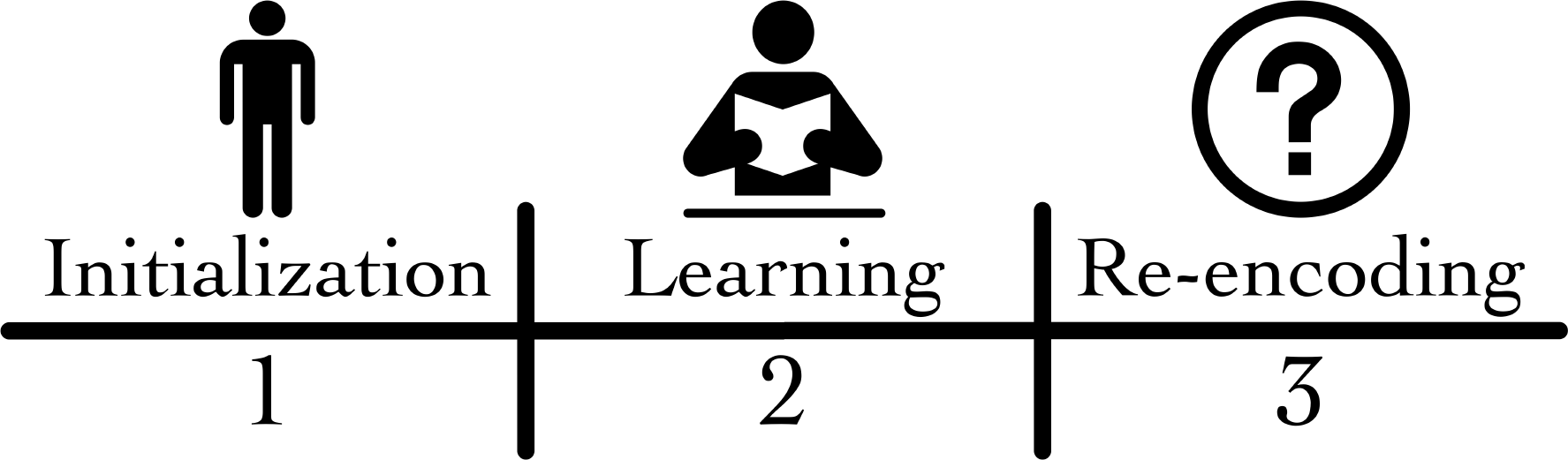
Paradigm time points. Model evaluation occurs at three time points in the RIF task. Time point 1 is at model initialization before any learning has occurred. Time point 2 is after the dataset has been learned but before re-encoding of the Rp+ pairs. Time point 3 is after Rp+ re-encoding has occurred.

## Results and Analysis

The primary objective of this work is to investigate the extent to which a model using BCM to instantiate the NMPH can produce RIF. With that said, solely evaluating model performance misses much of the neural and cognitive nuance that makes RIF a complex and scientifically worthwhile phenomenon. Beyond behavioral data, neuroimaging studies of RIF have demonstrated representational dynamics that both hint at underlying mechanisms (Wimber et al., 2015) and must be explained by a theoretical account of RIF, as by Ritvo et al., (2023). Therefore, these analyses are geared not just towards demonstrating non-monotonic plasticity’s sufficiency for RIF, but scrutinizing the model’s representational dynamics. Doing so will shed light on how non-monotonic plasticity produces RIF.

As a further note, much of the analyses were conducted over multiple runs of the same experiment to account for the inter-trial variability inherent to a model with random initializations. When discussing this work, as the specific number of trials is largely arbitrary, precise statistical measures such as p-values are likewise easily altered. To that end, the statistical significance—or lack thereof—of results is commented on but precise statistical values are not reported. Instead, emphasis is placed on the presence and validity of statistical differences.

### Memory performance

Both the non-monotonic and monotonic models’ performance on the paired associates recall task was assessed 100 times with the mean proportion of correctly recalled items plotted across the three time points. First, Rp+ memory performance followed an expected trend (Fig. 6a). For both the experimental and baseline model, the proportion of correctly recalled pairs increased from less than 40% at initialization to in excess of 80% following study. Performance increased again after Rp+ re-encoding to over 95%. The close correspondence between Rp+ performance for the two models was encouraging as it indicates the non-monotonic plasticity function does not impede the model’s ability to learn.

**Figure 6:**
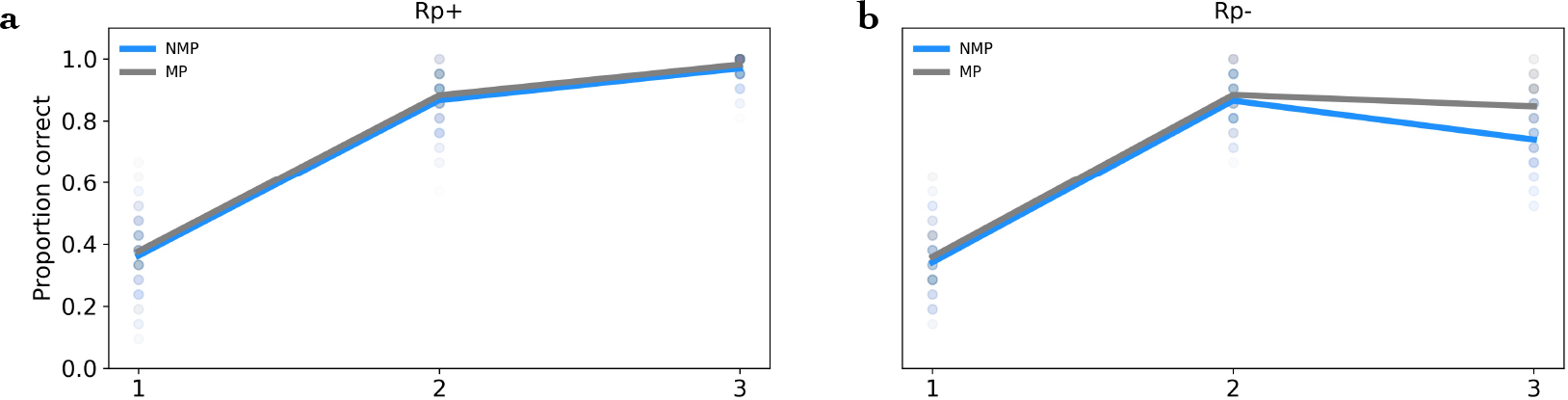
RIF performance. Mean performance for the non-monotonic (blue) and monotonic (gray) models, averaged across 100 runs. Lines are mean values and each dot is the result from a single run. The y axis is percent of pairs correctly recalled and the x axis are experimental time points. Rp+ sees consistent performance improvements as both models learn the dataset and practice the Rp+ pairs (**a**). By contrast, there’s an Rp-performance divergence between the two models after Rp+ re-encoding (**b**). While the non-monotonic model evidences a significant performance decline, consistent with RIF, the monotonic model does not.

Rp-performance, however, is the target of memory interference and primary metric of interest (Fig. 6b). Here, for the first two time points, there is a close correspondence between the two models. At the third, however, following Rp+ re-encoding, there is a divergence. Whereas the monotonic baseline model performs almost identically after Rp+ re-encoding as before, the non-monotonic experimental model displays a significant Rp-performance decrease, falling by approximately 20%. This constitutes the primary behavioral component of RIF: diminished memory performance on Rp-items as a result of practicing Rp+ items. As such, while the monotonic baseline leaves Rp-performance largely unchanged, evidencing no RIF, adding a non-monotonic domain to the plasticity function produces significant RIF.

### Representational divergence

Neuroimaging studies have shown that RIF not only involves changes in memory behavior but also in memory representation. As a consequence of Rp+ practice, the representations for Rp+ and Rp-items in memory were found to diverge, as measured by human fMRI (Wimber et al., 2015; Favila et al., 2016). To test this finding in the model, cosine similarity between Rp+ and Rp-hidden representations were calculated before and after Rp+ re-encoding, revealing the number of units shared between them. As with performance tests, 100 trials were run.

The non-monotonic model evidenced a significant decline in similarity, with mean similarity falling in excess of 10% (Fig. 7a). This finding is consistent both with neuroimaging studies showing representational divergence between Rp+ and Rp-in RIF (Wimber et al., 2015), and also with Ritvo et al. (2023), whose model similarly evidenced increased representational difference between two pairs upon one pair’s re-encoding. This is in stark contrast with the monotonic model which saw no significant change in representational similarity between Rp+ and Rp-after Rp+ re-encoding (Fig. 7b). The asymmetry of this representational shift was evaluated by taking the cosine similarity of Rp+ and Rp-representations before and after Rp+ re-encoding (Fig. 7c). Ritvo et al. (2023) found higher degrees of similarity for re-encoded pairs—Rp+ in our case—than for non re-encoded pairs— Rp-. We found much the same. On average, Rp+ representations before and after re-encoding shared over 2.5 out of the k=5 active hidden units. Rp-representations, by contrast, were largely unique, with most trials producing no shared units and no trials producing more than one shared unit. Rp-is thus the locus of the bulk of the representational divergence between Rp+ and Rp-. As a result of Rp+ re-encoding, Rp-representations are significantly altered whereas Rp+ representations are only moderately altered. In sum, this test of representational divergence found that, as a result of Rp+ re-encoding, the representations of Rp+ and Rp-patterns diverge. This divergence is predominantly driven by Rp-pairs as their representations differentiate significantly more than Rp+. It is thus clear that Rp+ re-encoding disrupts Rp-representations, and that, as representational divergence was not observed in the monotonic model, this disruption is driven by non-monotonic plasticity.

**Figure 7:**
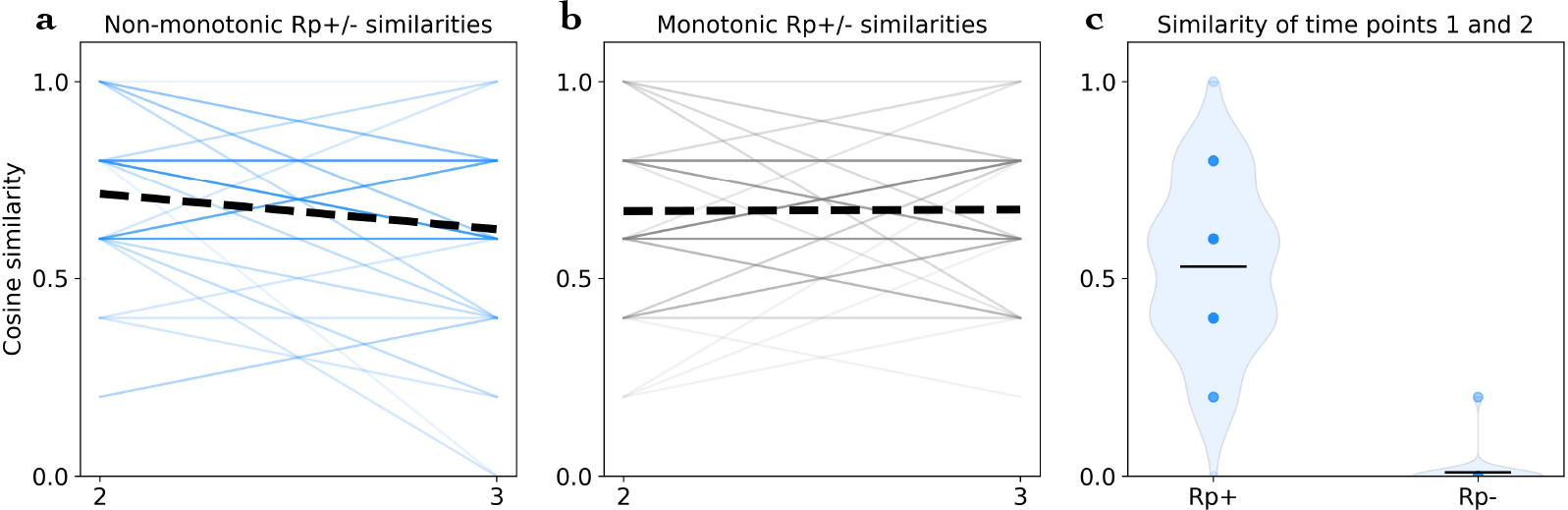
Representational divergence. The similarity between representations of Rp+ and Rp-pairs diverge significantly after Rp+ re-encoding—from time point two to time point three (**a**). Colored lines are each of 100 runs with the dashed line being the mean. Cosine similarity clusters for each of the k=5 units active in the hidden layer. There is no significant change in similarity between Rp+ and Rp-pairs for the monotonic baseline model (**b**). When evaluating the similarity of Rp+ and Rp-representations before re-encoding to Rp+ and Rp-representations after re-encoding (**c**), we found high levels of similarity for Rp+ pairs and low levels for Rp-. This indicates that Rp-pairs drive the representational divergence as re-encoding disrupts Rp-representations more than Rp+. Dots are individual runs and horizontal lines are means.

### Weight divergence is driven by non-monotonicity

To evaluate representational disruption as non-monotonic plasticity’s mechanism of RIF, we sought to link disruption to the plasticity function. As there is a precise manipulation between the non-monotonic and monotonic models, the locus of interference and the RIF characteristics it drives must be the lower portion of the plasticity function—the negative domain in the non-monotonic function that is bound to zero in the baseline model. Since this domain is the sole difference between the two models, it must be instrumental. The key question is, of the hidden units caught in this domain during Rp+ re-encoding, which subpopulation is responsible for the interference that gives rise to RIF and what impact does non-monotonicity have on those units’ weight updates?

To address these questions, the RIF task was halted at the second time point after the models have learned the dataset but before Rp+ re-encoding. A random Rp+ and Rp-pattern were then selected from the same category. Although designated Rp+ and Rp-, as the models have not undergone any further re-encoding, the Rp designation is irrelevant and the two patterns will be referred to as pair A and pair B. The two pairs were then presented to the model in turn, and the net input to each hidden unit and the matrix of weight updates each pair would induce were recorded. The model itself was kept frozen between pair presentations as the objective is to evaluate the weight changes each would hypothetically induce were it used for re-encoding.

The percentile rank of each hidden unit’s net input from pairs A and B were plotted against each other (Fig. 8a, b). As the pairs were drawn from the same category, the positive correlation is expected. The vector of hypothetical weight updates induced by pair A or pair B were then compared using cosine similarity. The similarity measures were then subtracted off one to convert them to a dissimilarity index and used to set the size of each point on an exponential scale. As such, the size of each hidden unit’s point corresponds to the dissimilarity between its weight updates for pairs A and B.

**Figure 8:**
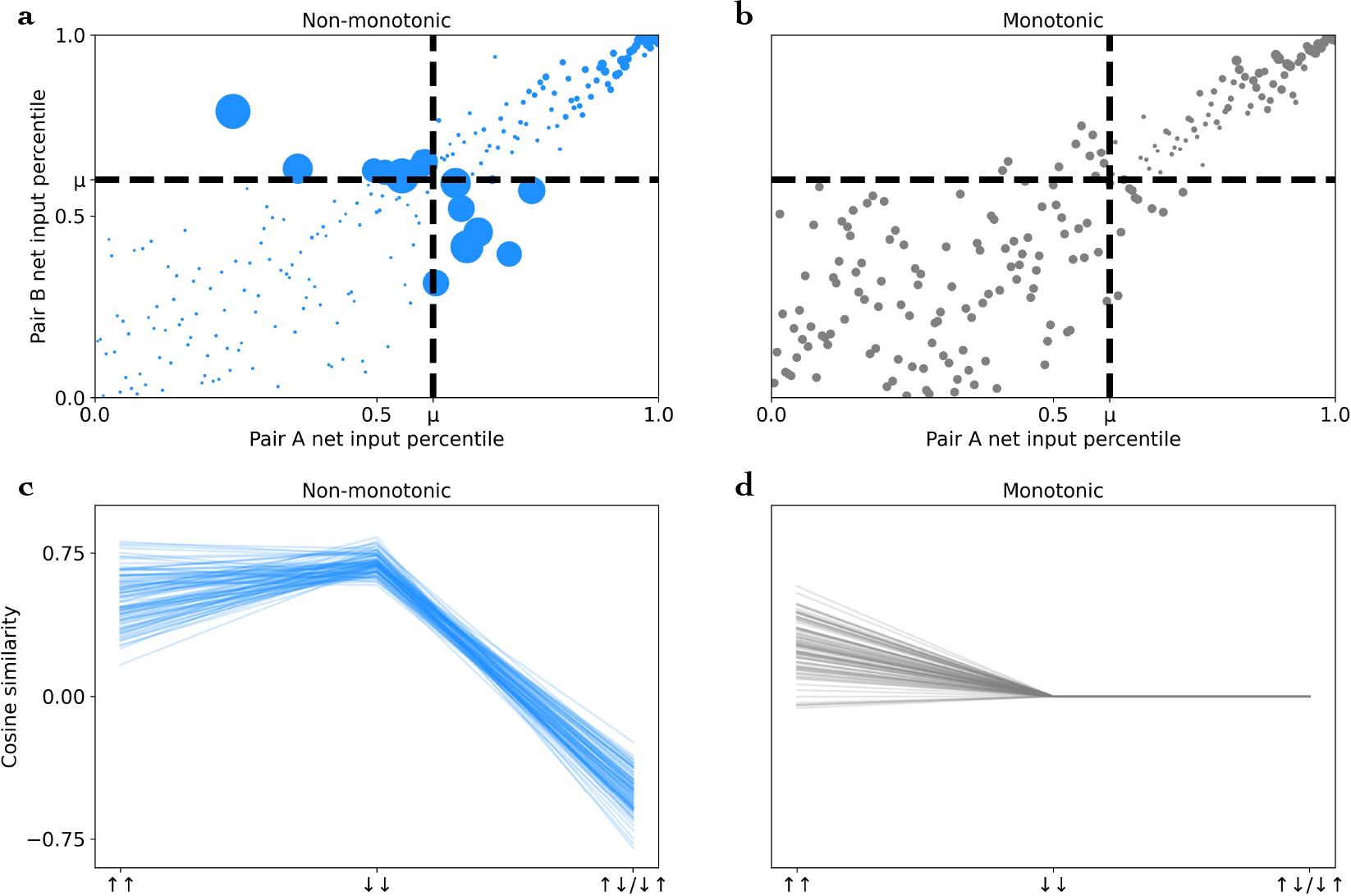
Non-monotonicity drives interference through diverging weight updates. Plots of the net input each hidden unit would receive from two randomly selected pairs of the same category, pair A and pair B, after learning the dataset but before Rp+ re-encoding (**a, b**). The size of each point is set by the dissimilarity between the weight updates incentivized by pairs A and B on an exponential scale for visualization purposes. The non-monotonic model shows that the hidden units most stimulated by one pair and least stimulated by the other (top left and bottom right quadrants) are the primary sources of divergence (**a**). By contrast, the monotonic model evidences consistent levels of weight update similarity throughout (**b**). Computing cosine similarity between weight updates for pairs A and B over 100 runs confirms that the hidden units falling in the same stimulation domain for both pairs—either consistently above *μ* or consistently below (⇈ and ⇊)—induce weight updates with a high degree of similarity (**c**). By contrast, the hidden units stimulated less than *μ* by one pair and more than *μ* by the other (⇅ / ⇵) are the source of diverging weight updates, and thus of inter-pattern interference (**c**). This highlights diverging weight updates as the mechanism behind representational divergence in the RIF task and the performance declines that accompany it. As the monotonic model doesn’t update the weights of its least active units, they undergo no change, meaning the model has no driver of representational divergence (**d**).

The resulting graph reveals an interesting trend. While the units that fall in the negative domain of the plasticity function for both pairs (**a**_**i**_ < *μ*)—the lower left quadrant—have weight updates that are minimally dissimilar between pairs A and B, the units that receive the least stimulation from one pair (**a**_**i**_ < *μ*) but the most from the other (**a**_**i**_ > *μ*)—those in the negative domain of the plasticity function for one pattern and the positive domain for the other—uniformly have highly dissimilar weight updates for pairs A and B (Fig. 8a). By contrast, all hidden units’ weight updates in the monotonic baseline model are consistently similar between the two patterns, with som slight variability observed for the highly stimulated units in the top right quadrant (Fig. 8b).

To reframe this trend, cosine similarities between the weight update matrices for pairs A and B were computed for three different populations of hidden units: for the units stimulated more than *μ* by both pairs (⇈), for the units stimulated less than *μ* by both pairs (⇊), and the units stimulated less than *μ* by one pair and more than *μ* by the other (⇅/ ⇵). As no re-encoding has occurred, the order of which pair produces stimulation greater than *μ* or less than *μ* is irrelevant so they are grouped together for analysis. These three categories correspond to the upper right, lower left, and upper left/lower right quadrants, respectively. Doing so confirms that the units in the (⇅/ ⇵) population are the driver of the pairs’ diverging weight updates, as the weight update similarities for these units are significantly lower than for all units in other quadrants (Fig. 8c). The monotonic baseline model, on account of its lack of a negative domain in its plasticity function, is unable to use the (⇅/ ⇵) population of units to drive divergence (Fig. 8d). Instead, as the least active units in the monotonic baseline model have their weight updates scaled by *ϕ*=0, these units undergo no weight change at all, producing no interference and explaining the lack of RIF.

These results indicate that the diverging representations of Rp+ and Rp-patterns, and the accompanying decline in Rp-recall performance, are the result of Rp+ re-encoding inducing weight updates that erode the model’s ability to reconstruct Rp-patterns. Such erosion is driven by units that fall in different domains of the plasticity function for the two pairs: the negative domain for Rp+ and the positive domain for Rp-. This explanation is consistent with the mechanics of the non-monotonic plasticity function. As the units falling in the negative domain of the function have their weights tuned away from the input pattern, the weight structure of these units is interfered with. Such interference is of little consequence for the practiced Rp+ pattern as the k-winner non-linearity ensures that only the most active units, whose weights are tuned towards the input, are used for pattern reconstruction. In essence, there is a guarantee that the units used to reconstruct the Rp+ pattern will not have their weight structure eroded by the negative domain of the non-monotonic plasticity function. Rp-, however, has no such guarantee. Some subset of the units used to drive Rp-reconstruction are liable to fall in the negative domain during Rp+ practice, and thus to have their weight structure disrupted. This explains both the Rp-performance decline that defines RIF and the asymmetrical representational divergence observed by Ritvo et al. (2023) and replicated in the prior section. This constitutes a mechanistic explanation of how non-monotonic plasticity implements RIF.

### Limitations

The model’s simplicity introduces several limitations. The human RIF literature is expansive and, besides the core RIF task, includes extensions that our model does not capture. Reverse RIF is an example. In reverse RIF, following Rp+ practice, the Rp-pairs are re-encoded. Doing so not only rescues Rp-memory performance but does so without impeding Rp+ performance. This is explained by increased representational difference between Rp+ and Rp-, with the point being RIF and subsequent reverse RIF see increasing degrees of representational differentiation—shifting patterns to forms that minimize the potential for interference without compromising performance (Hulbert & Norman, 2015; Ritvo et al., 2019).

The model’s ability to perform reverse RIF was evaluated by extending the RIF paradigm. A fourth time point was added during which the models practiced the Rp-pairs. Results were mixed. Although Rp-performance was rescued by reverse RIF, as predicted by the human subjects data (Fig. 9b), Rp+ performance was negatively impacted (Fig. 9a). These trends were largely mirrored in the monotonic baseline model, but, as that model did not evidence RIF, it cannot be evaluated for reverse RIF. The non-monotonic model therefore does not generalize to the reverse RIF task.

**Figure 9:**
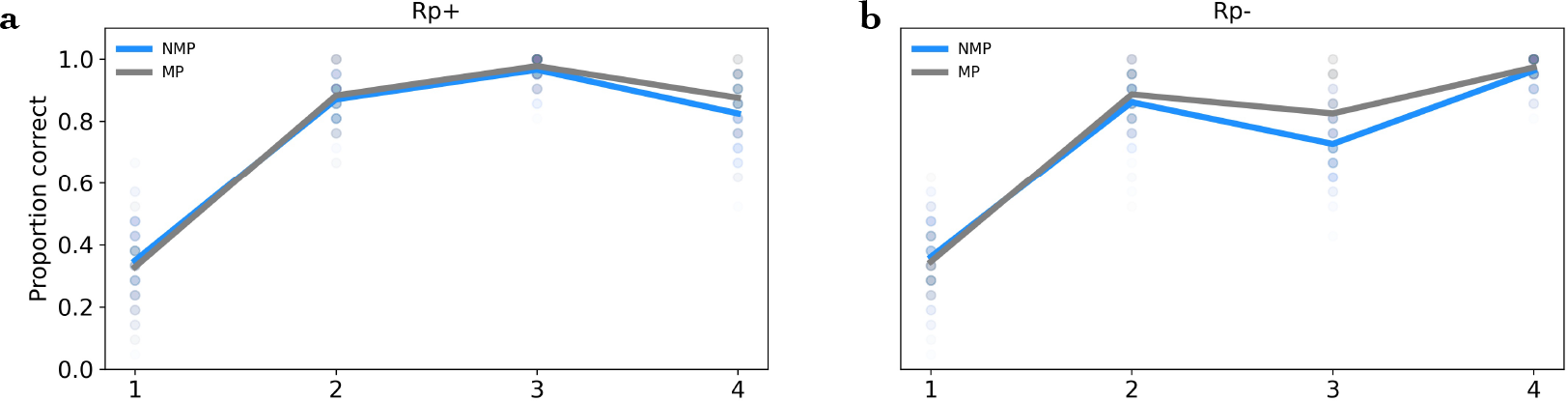
Reverse RIF failure indicates model limitations. Lines are mean performance of 100 runs of the reverse RIF task. Each dot is a recording from a single run. Even though Rp-performance is rescued (**b**), consistent with human subjects data, Rp+ performance declines for both models (**a**), contrary to human subjects experiments.

Several factors might explain the non-monotonic model’s inability to perform reverse RIF: the network may lack sufficient depth, the vector-based paired associates task may be ill-suited, or using a non-monotonic scalar of Hebbian learning may be a sub-optimal model of synaptic plasticity. Regardless, future work can develop more robust expansions of the proposed non-monotonic model to establish the sufficient mechanisms for a fuller suite of RIF-associated phenomena.

## Discussion

Here, we replicate Ritvo et al (2023) by demonstrating non-monotonic plasticity’s ability to bring about RIF in a neural network model. In sum, non-monotonic plasticity is shown to erode the model’s weight structure when re-encoding Rp+ pairs, precipitating inter-pattern interference that drives representations apart and causes the un-practiced Rp-pairs’ performance to decline. These are two primary characteristics of RIF. The ability of the non-monotonic model to capture them—and the monotonic baseline model’s inability—indicates that non-monotonicity is sufficient for these aspects of RIF. These RIF characteristics are produced by non-monotonic plasticity enforcing weight structure erosion of the units composing the Rp-reconstruction. Rp+ pairs are protected from this erosion as the units used for their reconstruction, by definition, are tuned towards their input pattern by the positive domain of the plasticity function, *ϕ*. This phenomenon explains the novel prediction found in Ritvo et al. (2023) that the representational divergence between pairs is driven by the unpracticed pair—in this case Rp-. The non-monotonic model’s ability to drive the behavioral and representational characteristics of RIF suggested it may likewise facilitate reverse RIF. However, we found the non-monotonic model was unable to perform reverse RIF.

Several promising directions of future research may shed light on the parameters of this limitation. One such direction are the PFC-HPC interactions that are believed to underpin goal-directed memory recall (Eichenbaum, 2017). A competing mechanistic theory of RIF, the active forgetting hypothesis (AFH; Anderson & Hulbert, 2021), centers on such interactions. Previous theoretical explorations of the NMPH and RIF have argued that PFC input plays a mediating but not a necessary role in RIF (Norman et al., 2007). Perhaps such additional inputs are required in reverse RIF. An expansion of this modeling work could therefore adopt this idea by using a dynamic goal-congruent filter to sparsify hidden layer activity, replacing the k-winner non-linearity. Doing so would emulate a key manifestation of PFC-HPC interactions and provide a parsimonious theory for how non-monotonic plasticity and PFC projections work in concert to produce a full suite of RIF properties. With that said, given the concise nature of the non-monotonic model, adding an analogue for PFC may not be necessary to capture reverse RIF. Instead, a more robust exploration of the model’s parameter space, of experimental paradigms, or of implementations of non-monotonic plasticity may be sufficient to capture reverse RIF. Several assumptions were built into the current model, such as a static value for in the non-monotonic plasticity function, that may significantly impact performance. In sum, while the current model has a robust but not all-encompassing ability to capture the full suite of RIF characteristics, expansions on this work may demonstrate that the gaps are filled by additional and complementary mechanisms, such as PFC projections, or more robust implementations of non-monotonic plasticity.

The consequence of this work has been to demonstrate that RIF is produced by a computational model of an idealized HPC that instantiates the NMPH; thereby replicating Ritvo et al. (2023) and demonstrating the robustness of non-monotonic plasticity as the computational mechanism of RIF. Our analyses found the model precipitated RIF through weight interference, driving apart the representations of competing items in memory. This effect similarly explains the novel prediction that representational divergence is predominantly driven by the unpracticed item. As these effects were not found in a monotonic control model, we thus provide grounding for non-monotonic plasticity’s sufficiency for RIF.

## Materials

All code used to generate the models and perform analyses will be made publicly available upon this manuscript’s publishing.

